# Accurate Prediction of Virus-Host Protein-Protein Interactions via a Siamese Neural Network Using Deep Protein Sequence Embeddings

**DOI:** 10.1101/2022.05.31.494170

**Authors:** Sumit Madan, Victoria Demina, Marcus Stapf, Oliver Ernst, Holger Fröhlich

## Abstract

Prediction and understanding of tissue-specific virus-host interactions have relevance for the development of novel therapeutic interventions strategies. In addition, virus-like particles (VLPs) open novel opportunities to deliver therapeutic compounds to targeted cell types and tissues. Given our incomplete knowledge of virus-host interactions on one hand and the cost and time associated with experimental procedures on the other, we here propose a novel deep learning approach to predict virus-host protein-protein interactions (PPIs). Our method (Siamese Tailored deep sequence Embedding of Proteins - STEP) is based on recent deep protein sequence embedding techniques, which we integrate into a Siamese neural network architecture. After evaluating the high prediction performance of STEP in comparison to an existing method, we apply it to two use cases, SARS-CoV-2 and John Cunningham polyomavirus (JCV), to predict virus protein to human host interactions. For the SARS-CoV-2 spike protein our method predicts an interaction with the sigma 2 receptor, which has been suggested as a drug target. As a second use case, we apply STEP to predict interactions of the JCV VP1 protein showing an enrichment of PPIs with neurotransmitters, which are known to function as an entry point of the virus into glial brain cells. In both cases we demonstrate how recent techniques from the field of Explainable AI (XAI) can be employed to identify those parts of a pair of sequences, which most likely contribute to the protein-protein interaction. Altogether our work highlights the potential of deep sequence embedding techniques originating from the field of natural language processing as well as XAI methods for the analysis of biological sequences. We have made our method publicly available via GitHub.

**The bigger picture:** Development of novel cell and tissue specific therapies requires a profound knowledge about protein-protein interactions (PPIs). Identifying these PPIs with experimental approaches such as biochemical assays or yeast two-hybrid screens is cumbersome, costly, and at the same time difficult to scale. Computational approaches can help to prioritize huge amounts of possible PPIs by learning from biological sequences plus already-known PPIs. In this work, we developed a novel approach (Siamese Tailored deep sequence Embedding of Proteins - STEP) that is based on recent deep protein sequence embedding techniques, which we integrate into a Siamese neural network architecture. We use this approach to train models by utilizing protein sequence information and known PPIs. After evaluating the high prediction performance of STEP in comparison to an existing method, we apply it to two use cases, SARS-CoV-2 and John Cunningham polyomavirus (JCV), to predict virus protein to human host interactions. Altogether our work highlights the potential of deep sequence embedding techniques originating from the field of natural language processing as well as Explainable AI methods for the analysis of biological sequence data.

**Highlights:**

- A novel deep learning approach (STEP) predicts virus protein to human host protein interactions based on recent deep protein sequence embedding and a Siamese neural network architecture
- Prediction of protein-protein interactions of the JCV VP1 protein and of the SARS-CoV-2 spike protein
- Identification of parts of sequences that most likely contribute to the protein-protein interaction using Explainable AI (XAI) techniques

**Data Science Maturity:** DSML 3: Development/Pre-production: Data science output has been rolled out/validated across multiple domains/problems

## Introduction

Viral infections can cause severe tissue-specific damages to human health. In case of the infection of brain cells severe neurological disorders can be the consequence (Swanson et al. 2015). Accordingly, prediction and understanding of tissue-specific virus-host interactions is important for designing targeted therapeutic intervention strategies. At the same time virus-like particles (VLPs), such as John Cunningham VLPs, open novel opportunities to deliver therapeutic compounds to targeted brain cells and tissues, because these proteins have the ability to cross the blood-brain barrier (Ye et al. 2021). Hence, it is also relevant from a therapeutic perspective to know the binding of VLPs to potential drug receptors in the brain.

The knowledge about virus-host interactions covered in databases like VirHostNet (Guirimand et al. 2015) is limited. While various experimental approaches exist to measure protein-protein interactions (PPI), including yeast two-hybrid screens, biochemical assays, and chromatography (Lalonde et al. 2008), these methods are often time consuming, laborious, costly, and difficult to scale to large numbers of possible PPIs. Thus, computational methods have been proposed that use various types of protein information to predict PPIs. Older approaches focused on predicting PPIs either using structure and/or genomic context of proteins (Skrabanek et al. 2008). Other approaches (Shen et al. 2007, Zhou et al. 2018) suggested classical machine learning algorithms (such as support vector machines) in combination with manually engineered features derived from protein sequences to predict PPIs.

In recent years, deep learning-based approaches (Sun et al. 2017, Wang et al. 2017, Xu et al., 2020, Tsukiyama et al., 2021) have become popular and have increasingly superseded traditional machine learning approaches for the prediction of PPIs. Often these approaches use known PPIs from established PPI databases (e.g., BioGrid, IntAct, STRING, HPRD, VirHostNet-Oughtred et al. 2020, Orchard et al. 2014, Szklarczyk et al. 2018, Keshava Prasad et al. 2009, Guirimand et al. 2015) to generate datasets to train deep neural network architectures. Some of these methods employ recent network representation learning techniques to complete a known virus-host protein-protein interaction graph (Du et al., 2021). Other authors focused on protein sequences to predict PPIs. For example, Sun et al. (2017) and Wang et al. (2017) proposed to employ a stacked autoencoder (SAE). Chen et al. (2019) developed a deep learning framework employing a Siamese neural architecture to predict binary and multi-class PPIs. Tsukiyama et al. (2021) recently proposed an LSTM-based model on top of a classical word2vec embedding of sequences to predict human-virus PPIs by using protein sequences. Using the same embedding technique, Liu-Wei et al. (2021) developed an approach, which predicts host-virus protein-protein interactions for multiple viruses considering their taxonomic relationships.

In the last few years, transfer learning-based approaches from the natural language processing (NLP) area have massively impacted the field of protein bioinformatics (Elnaggar et al. 2021; Min et al. 2019; Heinzinger et al. 2019). These methods are trained on a huge amount of protein sequences to learn informative features of protein sequences. For instance, Elnaggar et al. (2021) employed 2.1 billion protein sequences for the pre-training of ProtTrans, a collection of transformer models originally stemming from the NLP field. Such methods allow the transformation of a protein sequence into a vector representation, which can subsequently be used efficiently for various downstream tasks, e.g. protein family classification (Nambiar et al. 2020). There are several advantages of using the available pre-trained transformer models, such as avoiding the error-prone design of hand-crafted features to encode protein sequences and correspondingly a much more efficient development of new AI models with potentially higher prediction performance.

In this paper, we introduce a novel deep learning architecture combining the recently published ProtBERT (Elnaggar et al. 2021) deep sequence embedding approach with a Siamese neural network to predict PPIs by utilizing the primary sequences of protein pairs. While recent publications generally follow a similar strategy, they have employed more traditional sequence embedding methods (Tsukiyama et al. 2021). To our knowledge, our work thus constitutes the first attempt to evaluate the usage of most recent, pre-trained transformer models to obtain a deep learning-based biological sequence embedding for PPI prediction. After evaluating the promising prediction performance of our method (Siamese Tailored deep sequence Embedding of Proteins - STEP), we employ it for two use cases: i) predicting interactions of the John Cunningham polyomavirus (JCV) major capsid protein VP1 (UniProt:P03089) with human receptors in the brain, and ii) predicting interactions of the SARS-CoV-2 spike glycoprotein (UniProt:P0DTC2) with human receptors. Predicted interactions in both cases demonstrate a clear interpretation in the light of existing literature knowledge, hence supporting the biological relevance of predictions made by our method.

With this study, we make four contributions to the state-of-the-art. Firstly, we construct a novel deep learning architecture STEP for virus-host PPI prediction that requires only the protein sequences as the input and discards the need of handcrafted or other types of features. Secondly, we demonstrate that utilizing transformer-based models for PPI prediction achieves at least state-of-the-art performance for PPI prediction. In computer vision and NLP, such transformer-based models have shown that they are well suited for learning contextual relationships hidden in sequential data. However, these have not yet been applied to the field of PPI prediction. Hence, we use and build on the huge effort of Elnaggar et al. (2021), who published a pre-trained ProtBERT model that was trained on over two billion amino acid sequences. In addition, we demonstrate that using transfer learning in STEP achieves state-of-the-art performance, for which we evaluated STEP on multiple publicly-available virus-host and host-host PPI datasets. Thirdly, we predict interactions for two viruses that are known to cause serious diseases and provide an interpretation on those predictions demonstrating the support through existing literature knowledge. Lastly, we show how experimental XAI techniques could be used to identify regions in protein amino acid sequences that attribute to the prediction of protein-protein interaction.

## Experimental procedures

### Resource availability

#### Lead contact

Further information and requests for code and data should be directed to and will be fulfilled by the lead contact, Holger Fröhlich.

#### Materials availability

This study did not generate any physical materials.

#### Data and code availability

The data and source code is available at https://github.com/SCAI-BIO/STEP.

#### Construction of Datasets

#### Primary Data Sources

The following primary resources were employed to create training and test datasets in this work:

1. UniProt protein sequence dataset (The UniProt Consortium 2021) containing human protein sequences
2. UniProt mapping dataset (The UniProt Consortium 2021) containing mappings to other databases
3. VirHostNet dataset (Guirimand et al. 2015) including virus-host interactions of SARS-CoV-2 spike glycoprotein.
4. PPT-Ohmnet dataset (Zitnik et al. 2018) (https://snap.stanford.edu/biodata/datasets/10013/10013-PPT-Ohmnet.html) containing brain tissue-specific protein-protein-interactions
5. The Gene Ontology (GO) (Gene Ontology Consortium 2004) receptor protein dataset containing annotation of proteins as receptors and parts of protein complexes
6. Sequences of JCV major capsid protein VP1 (https://www.uniprot.org/uniprot/P03089, accessed on 18 November 2021) and SARS-CoV-2 spike glycoprotein (https://www.uniprot.org/uniprot/P0DTC2, accessed on 18 November 2021)
7. Pathogen-host PPI training and test set provided by Tsukiyama et al. (2021) (http://kurata35.bio.kyutech.ac.jp/LSTM-PHV/download_page, accessed on 18 November 2021) (used for comparative analysis)
8. Yeast PPI dataset from Guo et al. (2008) (used for comparative analysis)
9. Human PPI dataset from Sun et al. (2017) (used for comparative analysis)
10. Protein-protein interaction type prediction dataset SHS27k from Chen et al. (2019) (used for comparative analysis)
11. PPI binding affinity estimation dataset from Chen et al. (2019) (used for comparative analysis)

#### Construction of Brain-specific Protein-Protein Interactome Dataset

We chose the PPT-Ohmnet database (Zitnik et al. 2018) that includes tissue-specific human protein-protein interactions collected from various sources. PPT-Ohmnet only takes physical PPIs into account that are supported by experimental evidence (https://snap.stanford.edu/biodata/datasets/10013/10013-PPT-Ohmnet.html). More specifically, interactions contained in PPT-Ohmnet were collected from various curated databases such as TRANSFAC, IntAct, MINT (Menche et al. 2015). The tissue information for an interaction was inferred through the low-throughput tissue-specific gene expression data (Greene et al. 2015). The protein-protein interactome can be considered as a graph, in which the proteins represent nodes and the interactions between them are considered as edges. Furthermore, every edge contains the information about the tissue type. In total, there are 144 tissue types with 4,510 proteins (nodes) and about 3,666,563 non-unique edges (interactions) in the whole PPT-Ohmnet graph. More details about the creation and content of the PPT-Ohmnet database can be found in (Menche et al. 2015, Greene et al. 2015).

We extracted all tissue types and manually filtered the ones specific for the brain. In total, 36 brain-specific tissue types could be found from a total of 144 in the PPT-Ohmnet database (Figure S1). Using the information about brain tissue specific co-expression of proteins, we filtered the PPT-Ohmnet interactome. The final brain tissue-specific interactome contains 3,548 proteins (nodes) and 977,990 non-unique edges (interactions). Furthermore, the interactome contains 56,021 unique edges, from which 1,466 PPIs that interact with themselves were excluded. In total, 54,555 PPIs were used for further analysis. Supplementary Figure S1 shows the distribution of proteins and their interactions for each brain-specific tissue type. File S1 contains the brain-specific tissue types.

We further enriched each interaction with information about employed experimental detection methods. This information is not included in PPT-Ohmnet, hence, we used BioGRID and IntAct as the two largest PPI databases to extract the experimental procedures, such as “pull down”, “two hybrid”, by which the interactions were originally discovered. The list of experimental procedures was further manually curated to filter out detection methods considered as unreliable. Only PPIs detected by methods considered as reliable were used for further processing.

To train deep learning models we retrieved the sequences of all proteins in our PPIs from the UniProt database. We downloaded the human proteins dataset from the manually curated part of UniProt – so-called SwissProt (The UniProt Consortium 2021). Next, we extracted for all proteins their sequences and metadata such as name, ID, label. In total, sequences for 20,396 human proteins could be found. Finally, we filtered the PPIs and human receptor proteins for which we found the sequences.

#### Construction of SARS-CoV-2 Protein-Protein Interactome Dataset

As a second dataset, we used the VirHostNet database (Guirimand et al. 2015) to collect all PPIs between SARS-CoV-2 and human proteins. We extracted for all human and SARS-CoV-2 proteins their sequences and metadata such as name, ID, label from SwissProt. Our VirHostNet interactome contained 334 PPIs involving 338 proteins between SARS-CoV-2 and Homo sapiens.

#### Collection of Human Receptor Proteins

To extract human receptor proteins, we first performed a search in GO for the term “receptor”. The GO branch annotation “Cellular components” was used to filter only for proteins. The GO annotation “organism” was used to filter for human proteins. In total 2075 results were found, in which 2,059 human receptor proteins and 16 human protein complexes were included. For further analyses, we only focused on human receptor proteins, for which we retrieved associated protein sequences from SwissProt. In total, sequences for 2,027 human receptor proteins could be found. Supplementary File S2 includes the list of identified human receptor proteins.

#### Preparation for Positive Unlabeled Learning

The goal of PPI detection is to learn a model that is able to detect whether there exists an interaction between two proteins. This task is often considered as a binary classification problem that can be solved by training a classifier to distinguish between positive and negative instances. However, the available PPI databases just contain positive, true interactions. Interactions not listed in a PPI database might still exist, but are possibly unknown today. Positive unlabeled (PU) learning is a scheme where a machine learning algorithm only has access to positive and unlabeled instances (Bekker and Davis 2020, Sansone et al. 2018). In PU learning all non-existing or unknown PPIs can be considered as “unlabeled” or as “pseudo-negatives”, however, they might also contain an unknown fraction of positive instances. Therefore, PU learning amounts to constructing a binary classifier that ranks instances with respect to the positive class conditional probability.

A popular strategy of PU learning is to first focus on selection of reliable negative instances. In a second step then a conventional binary classifier is trained on positive and selected negative instances (Bekker and Davis 2020). There are two types of strategies to sample pseudo-negative instances: random sampling or similarity-based sampling. With the random sampling strategy, the negative instances are created by randomly exchanging one of the partners in an interaction protein pair. While the similarity-based sampling considers the sequence similarity (or dissimilarity) of proteins. An example of this strategy is the Dissimilarity-Random-Sampling method (Eid et al. 2015), also used by Tsukiyama et al. (2021), which follows the hypothesis, if two viral proteins have similar sequences, a human protein that interacts with one of them cannot be paired with the other as a negative example. Sampling of highly dissimilar negative samples might result in overoptimistic classification performances (Tsukiyama et al. 2021). Therefore, in our work, we applied the random sampling approach to create negative instances. A major challenge in this context is the high class imbalance between positive and unlabeled training instances in our data. Hence, we decided to randomly subsample an equal number of pseudo-negatives.

### Architecture and Transfer Learning of STEP

We employ a deep Siamese neural network architecture while using transfer learning to learn relevant, latent features of PPI pairs based on protein sequences.

#### ProtBERT: Pre-trained Embeddings of Protein Sequences

ProtBERT (Elnaggar et al. 2021) is a pre-trained model trained on around two billion protein sequences using a masked language modeling objective (Devlin et al. 2018). It is based on the BERT model (Devlin et al. 2018) that was developed for the natural language domain. Hereby, ProtBERT considers protein sequences as sentences and the so-called building blocks of proteins - amino acids - as vocabulary. The ProtBERT model, specifically the BFD variant (Elnaggar et al. 2020) used in this work, consists of 30 layers with 16 attention heads and 1024 hidden layers. It was trained by using the Lamb (You et al. 2019) optimizer for around 23.5 days on 128 compute nodes each containing 1024 Tensor Processing Units (TPUs). During training, the language model learns to extract the biophysical characteristics of proteins from billions of protein sequences.

#### Siamese Neural Network Architecture

Given a pair of proteins, we first obtained their sequences. These sequences were then fed into a Siamese model architecture (Figure 1), in which the pre-trained ProtBERT model was used to obtain embeddings of both protein sequences. There are various ways to infer the relation between sequence embeddings. Some researchers focus on concatenation and others focus on element-wise multiplication (also known as Hadamard product) of both sequence embeddings. In this work we implemented an integration layer that uses the Hadamard product to combine the sequence embeddings as it is often found to be the most effective way to model symmetric characteristics of proteins (Chen et al. 2019).

**Figure 1.**
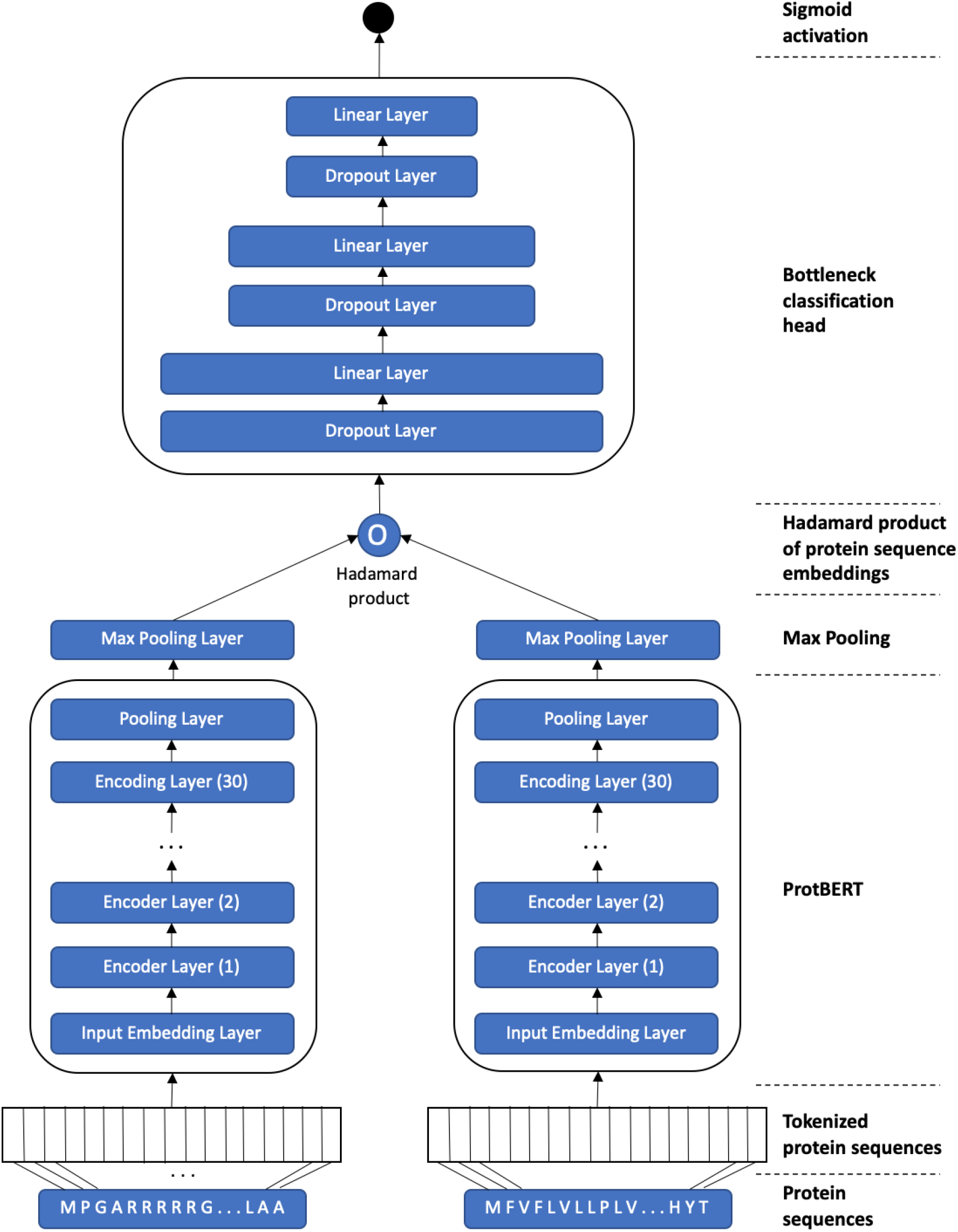
Architecture of our STEP model that employs the Siamese neural network while using the ProtBERT embeddings.

#### Classification Head for PU Learning

On top of the integration layer, we added a classification head represented by multiple hidden layers (Figure 1). We designed the classification head as a bottleneck-shaped architecture with a combination of dropout and linear layers, which ended in an output layer using a logistic function and thus allowed to rank protein pairs as either more likely to interact (positive) or not (negative). Notably, a network with bottleneck structure introduces a gradual decrease of the number of neurons per layer that allows the network to focus on relevant information and discards redundant or irrelevant information.

#### Evaluation Criteria

We evaluated our models using an independent test data set. This consisted of a defined fraction of known PPIs taken at random and excluded from training plus a specified fraction of pseudo-negatives that were not part of the training set. The performance was measured using the Area Under Receiver Operator Characteristic Curve (AUC) and Area Under Precision Recall (AUPR).

It should be re-emphasized that in our data negative samples are those protein pairs, for which an interaction is unknown. Therefore, we evaluated the ability of our models to enrich true positives at the beginning of a predicted ranking of potential PPIs. This ability is exactly reflected by AUC and AUPR measures, which are thus frequently employed in the literature about PU learning (Sansone et al. 2019). Notably, from a theoretical point of view the AUC estimated via PU learning and the one from a fully-labeled dataset are proofably linearly correlated (Menon et al. 2015).

#### Hyperparameter Optimization

To tune our system, we performed an extensive Bayesian hyperparameter optimization (Bergstra et al. 2013) using the training data. Due to the huge amount of training time for a single trial, hyperparameter candidates were evaluated using a single validation set consisting of a specified fraction of known PPIs plus an equal amount of sub-sampled negatives. For each trial, intermediate and final performances were assessed using the area under the receiver operating characteristic curve (AUC) measure and captured in an SQL database for later analyses. The captured data was also used by the pruning process of Optuna to stop the unpromising trials at an early stage (Akiba et al. 2019). Each optimization trial was executed on 2x A100 NVIDIA GPUs with VMEM of 32GB and 5 trials were executed parallely by employing 10x GPUs in total. The whole optimization process took 10 full days by executing 116 trials in total. The evaluated hyperparameter ranges and the best parameters are illustrated in Supplementary Table S1 and S3.

### Making STEP Models Explainable: Analysis of Integrated Gradients

One of the main criticisms of modern deep learning approaches is their often perceived black box character. To address this concern we aimed to understand the influence of individual amino acids on model predictions. For that purpose, we used the integrated gradients method (Sundararajan et al. 2017), which offers an intuitive and mathematically sound approach to explain predictions made by a deep neural network. Integrated gradients require no modifications to the trained model. Given an input sample (*x* ∈ *R*^*n*^), integrated gradients rely on a baseline / reference input sample (*x′* ∈ *R*^*n*^), which we constructed using the concatenation of one class, multiple padding, and one separator token. For a STEP model *F* : *R*^*n*^ → [0,1], integrated gradients are then obtained by accumulating the partial derivatives 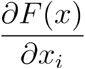 with respect to input feature *i* while moving from the reference to *x*′ the actually observed input *x* (Sundararajan et al. 2017):

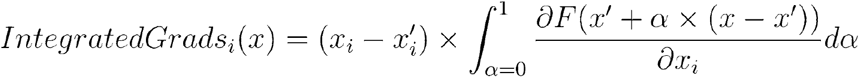

We used 1000 steps to approximate the integrated gradients, as suggested by Sundararajan et al. (2017) for highly nonlinear networks.

### Gene Set Enrichment Analysis (GSEA)

To better understand the biology of all ranked predictions in the individual use cases, we performed a GSEA to investigate an enrichment of gene sets listed in the Molecular Signatures Database (MSigDB; Subramanian et al., 2005). We downloaded molecular function gene sets of The Gene Ontology (GO) included as the collection C5 from MSigDB (v7.4, MSigDB/c5.go.mf.v7.4.symbols.gmt and MSigDB/c5.go.bp.v7.4.symbols.gmt). We considered a GO term to be statistically significant if, after applying the multiple hypothesis testing correction with the Benjamini–Hochberg method (Benjamini and Hochberg, 1995), its adjusted p-value was below 0.01.

## Results

### Comparative Evaluation of STEP with State-of-the-art Work

We performed a head-to-head comparison of our STEP architecture on three different datasets published by Tsukiyama et al. (2021), Guo et al. (2008), and Sun et al. (2017). Tsukiyama et al. (2021) recently published the LSTM-PHV Siamese model, which employs a more traditional word2vec sequence embedding. The dataset published by the authors consists of host-virus PPIs that were retrieved through the Host-Pathogen Interaction Database 3.0 (Ammari et al. 2016). In total, the dataset consists of 22,383 PPIs with 5,882 human and 996 virus proteins. Additionally, it includes artificially sampled negative instances with the positive to negative ratio of 1:10. The authors themselves compared LSTM-PHV on their dataset against a Random Forest approach by Yang et al. (2020). Guo et al. (2008) published a Yeast PPI dataset and used support vector machines to build a PPI detection model. Sun et al. (2017) created a dataset using human protein references database (HPRD), which contains human-human PPIs. Tsukiyama et al. (2021) and Guo et al. (2008) performed a 5-fold cross validation (CV) experiment, whereas Sun et al. (2017) used a 10-fold CV setting. We evaluated our STEP architecture using the exact same datasets with the exact same data splits as the authors of the compared methods. STEP was initialized with the hyperparameters shown in Table S1). Table 1 shows the results of all experiments, demonstrating at least state-of-the-art performance of our method. Additionally, we can conclude that our approach compared on exactly the same data published by Tsukiyama et al. (2021) performs similar to their LSTM-PHV method and better than the approach by Yang et al (2020).

**Table 1:** Overview of the results of comparative evaluation of STEP on LSTM-PHV (Tsukiyama et al. 2021), Yeast (Guo et al. 2008), and Human PPI (Sun et al. 2017) datasets. For LSTM-PHV and Yeast PPI datasets, we applied a 5-fold cross validation similar to the authors of the given studies. For the Human PPI dataset of Sun et al. (2017), we applied a 10-fold cross validation for training the STEP models. The highest values are highlighted in bold. More details of each experiment can be found in Supplementary Tables S1, S2, and S3. NA = not available in original publication.

| **<u>Comparative analysis on host-virus PPI dataset from Tsukiyama et al. (2021) via 5-fold cross-validation</u>** |  |  |  |  |
| --- | --- | --- | --- | --- |
|  | **AUC** | **AUPR** | **F1** | **MCC** |
| **Tsukiyama et al. (2021)** | 97.58%<br>(+/-0.13%) | 93.86%<br>(+/-0.35%) | 91.00%<br>(+/-0.53%) | 90.30%<br>(+/-0.53%) |
| **STEP (ours)** | **98.72%**<br>(+/-0.16%) | **95.71%**<br>(+/-0.51%) | **91.53%**<br>(+/- 0.65%) | **90.82%**<br>(+/-0.72%) |
| **<u>Comparative analysis on single independent host-virus PPI test dataset from Tsukiyama et al. (2021)</u>** |  |  |  |  |
|  | **AUC** | **AUPR** | **F1** | **MCC** |
| **Yang et al. (2020)** | 96.3% | 81.0% | 72.4% | 69.7% |
| **Tsukiyama et al. (2021)** | 97.3% | 93.8% | **91.1%** | **90.4%** |
| **STEP (ours)** | **98.50%** | **94.50%** | 89.69% | 88.76% |
| **<u>Comparative analysis on Yeast PPI dataset from Guo et al. (2008) via 5-fold cross-validation</u>** |  |  |  |  |
|  | **AUC** | **AUPR** | **F1** | **MCC** |
| **Guo et al. (2008)** | NA | NA | 87.34%<br>(+/-1.33) | 75.09%<br>(+/-2.51%) |
| **Chen et al. (2019)** | NA | NA | 97.09%<br>(+/-0.23%) | 94.17%<br>(+/-0.48%) |
| **STEP (ours)** | 99.61%<br>(+/-0.10%) | 99.58%<br>(+/-0.17%) | **97.37%**<br>(+/-0.27%) | **94.77%**<br>(+/-0.54%) |
| **<u>Comparative analysis on Human PPI dataset from Sun et al. (2017) via 10-fold cross-validation</u>** |  |  |  |  |
|  | **AUC** | **AUPR** | **F1** | **MCC** |
| **Sun et al. (2017)** | NA | NA | 97.15% | NA |
| **STEP (ours)** | 99.74%<br>(+/-0.03%) | 99.66%<br>(+/-0.04%) | **98.84%**<br>(+/-0.09%) | 97.67%<br>(+/-0.18%) |

Finally, we also evaluated our STEP architecture on two additional tasks, namely PPI type prediction and a PPI binding affinity estimation using the data by Chen et al. (2019). For both tasks, we reached at least state-of-the-art performances with our approach (see Supplementary Material; Table S4).

### Prediction of JCV Major Capsid Protein VP1 Interactions

We split the brain tissue-specific interactome dataset including all positive and pseudo-negative interactions into training (60%), validation (20%), and test (20%) datasets. The validation set was used for tuning hyper-parameters of the model only (see Table S5). After tuning on the validation set, we used our best model to make predictions on the hold-out test set. Figure 2 illustrates the area under receiver operator characteristic curve (AUC) and Precision-Recall Curve (AUPR). The model achieved an AUC and AUPR of 88.78% and 88.32% on the unseen test set, respectively. Also on an extended test set with a ratio 1:10 of positive to pseudo-negative samples the results are quite stable (see Table S6).

**Figure 2.**
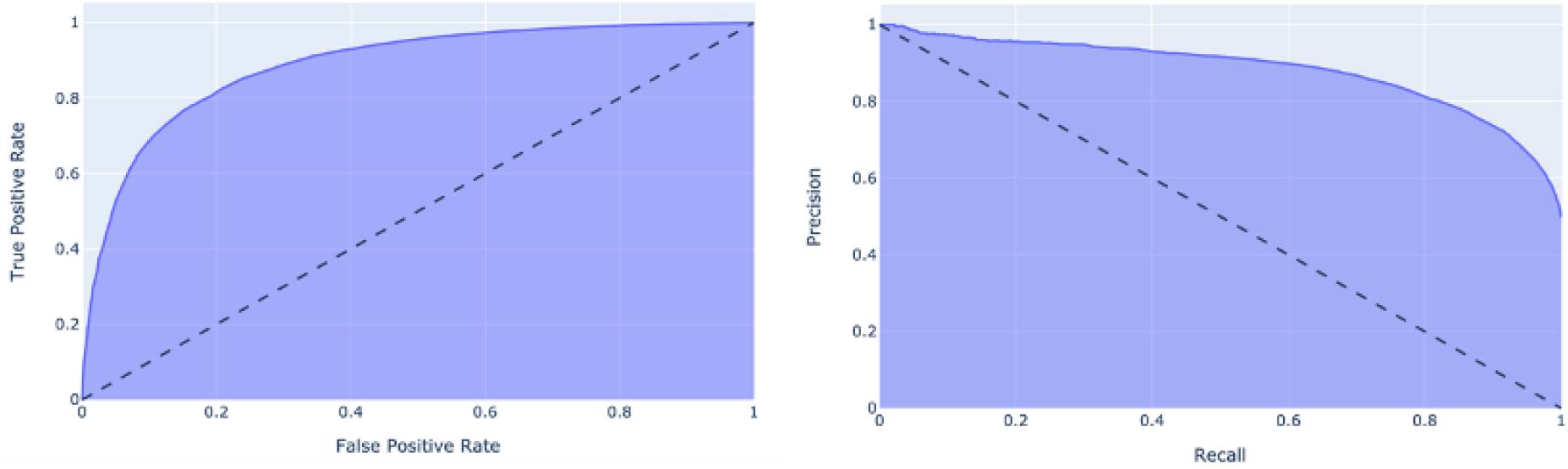
Receiver Operator Characteristic (ROC) curve (left) and precision-recall curve (right) obtained by applying the STEP-Brain model on unseen test data.

We used this STEP-Brain model to predict interactions of the JCV major capsid protein VP1 with all human receptors. Table 2 shows the top 10 predicted interactions that are ranked by the score retrieved by the logistic output function of the model. Supplementary File S3 contains all the predicted interactions. According to the method of integrated gradients, large parts of the VP1 sequence contribute to our model’s prediction of the protein-protein interaction with the top ranked receptor KIAA1549 (Figure S4). More specifically, signal peptide N-regions in KIAA1549 negatively contribute to the predicted class, whereas the beginning of the non-cytoplasmic domain region is contributing positively.

**Table 2:**
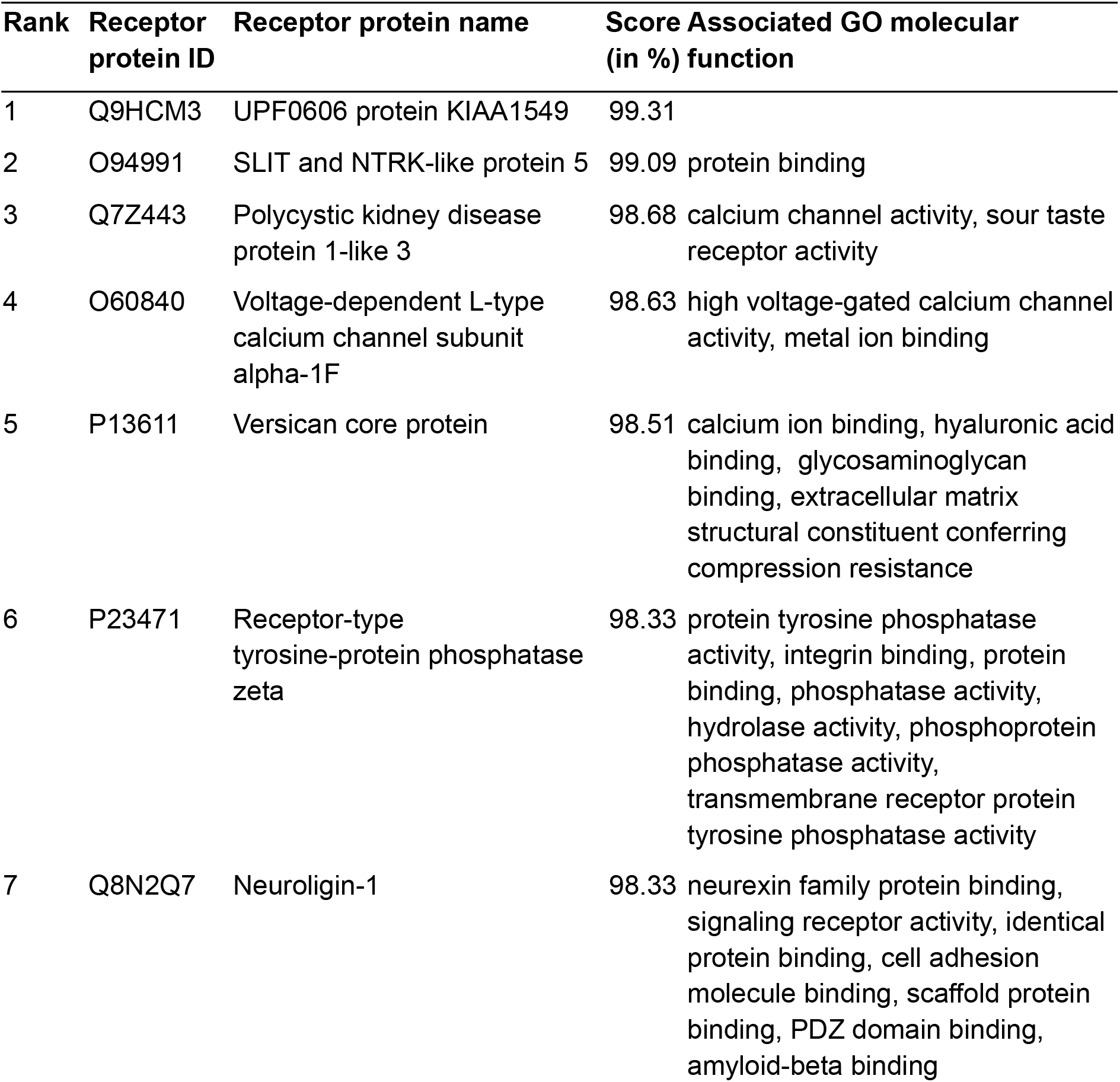
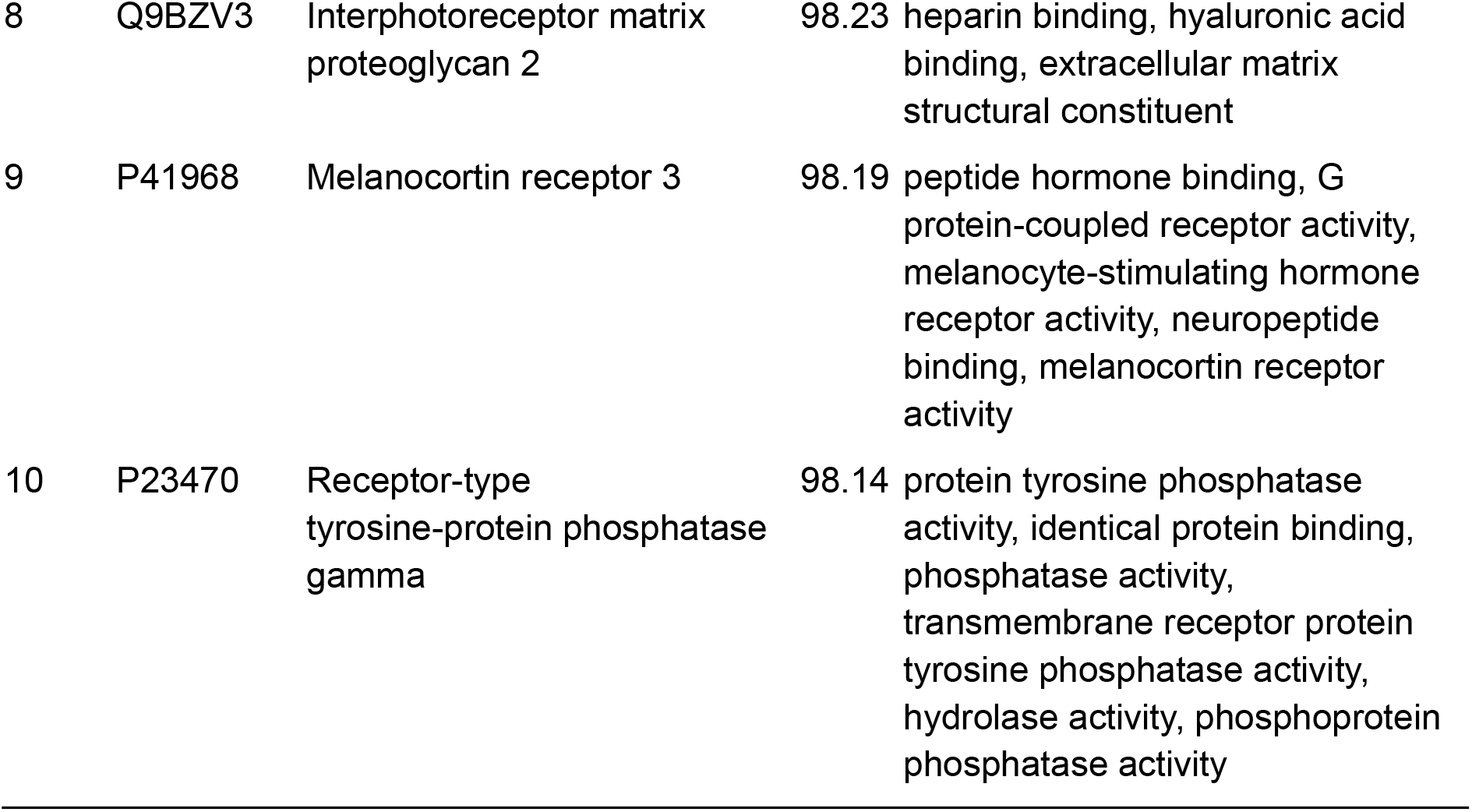
Top 10 predicted interactions of the JCV major capsid protein VP1 and human receptors ranked by the probability obtained by our model.

Altogether, we observed a strong enrichment of VP1 interactions predicted with olfactory, serotonin, amine, taste, and acetylcholine receptors (Figure S2). Notably, neurotransmitter (and specifically serotonin) receptors have previously been suggested to be the entry of the virus into myelin producing glial brain cells (Ferenczy et al. 2012), causing progressive multifocal leukoencephalopathy as a fastly progressing and life-threatening neurodegenerative disorder (Boothpur and Brennan 2010). Furthermore, we found an enrichment of tyrosine kinase activity, which is in line with the fact that tyrosine kinase inhibitors have been suggested as therapy against JCV (Querbes et al. 2004, Bennet et al. 2018).

We further performed an enrichment analysis with InterPro (Blum et al. 2021) protein domains for the predicted interactions between JCV major capsid protein VP1 and human receptors (Figure S5, Table S7). In line with the GO enrichment analysis, the two top-ranked protein domains InterPro:IPR006029 and InterPro:IPR006202 are neurotransmitter-gated ion channel transmembrane domains that open transiently upon binding of specific ligands, which then allow transmission of signals at chemical synapses (Kofuji et al. 1991, Wagner et al. 1991). Furthermore, the receptor-type tyrosine-protein phosphatase / carbonic anhydrase domain is enriched, which is in line with the enrichment of tyrosine kinase activity found via GO analysis. The enriched domains InterPro:IPR013106 (Immunoglobulin V-set domain) and InterPro:IPR007110 (Immunoglobulin-like domain) are both immunoglobulin-like domains that are involved in cell-cell recognition, cell-surface receptors and immune system response (Teichman et al. 2000), which play a role in the recognition of a virus protein.

### Prediction of SARS-CoV-2 Spike Glycoprotein Interactions

We performed a nested cross-validation procedure on the given SARS-CoV-2 interactions dataset. We used 5 outer and 5 inner loops to validate the generalization performance and while performing the hyperparameter optimization in the inner loop. In each outer run, we created a stratified split of the interactome into train (4/5) and test (1/5) datasets. In the nested run, we further split the outer train dataset into train (1/5) and validation (1/5) datasets, which were used to optimize the hyperparameters of the model using the respective training data. The performance of the classifiers was evaluated with AUC and was averaged over all nested runs. The best identified hyperparameters (see Table S8) were used to train the models in the outer loop. We retrieved a final generalization performance of 83.42% (+/-3.91%) AUC and 84.02% (+/-4.58%) AUPR that was calculated by averaging the prediction results of the outer loop (see Table 3).

**Table 3:** Results of the outer loop folds retrieved during the nested cross validation of STEP-Virus-Host model by using the test set with a ratio of 1:1 positive to pseudo-negative instances.

| Outer fold | AUC | AUPR |
| --- | --- | --- |
| 1 | 88.17% | 89.93% |
| 2 | 86.83% | 88.62% |
| 3 | 77.03% | 77.73% |
| 4 | 82.52% | 81.67% |
| 5 | 82.56% | 82.15% |
| **Mean** | 83.42% (+/- 3.91%) | 84.02% (+/- 4.58%) |

We used the STEP-Virus-Host model obtained from the best outer fold to predict interactions of the SARS-CoV-2 spike protein (alpha, delta and omicron variants) with all human receptors, which were not already contained in VirHostNet (see Supplementary Table S11, S12, S13). Supplementary File S4 contains all the predicted interactions for the omicron variant. Interestingly, for all virus variants the sigma intracellular receptor 2 (GeneCards:TMEM97; UniProt:Q5BJF2) was the only one predicted with an outstanding high probability (of >70% in all cases, Tables S11 -S13). Sigma 1 and 2 receptors are thought to play a role in regulating cell survival, morphology, and differentiation (Huang et al. 2014, Guo and Zhen 2015). In addition, sigma receptors have been proposed to be involved in the neuronal transmission of SARS-CoV-2 (Yesilkaya et al. 2020). They have been suggested as a target for therapeutic intervention (Abate et al. 2020, Gordon et al. 2020, Ostrov et al. 2021). Our results suggest that the observed antiviral effect observed in cell lines treated with sigma receptor binding ligands might be due to a modulated binding of the spike protein, hence inhibiting virus entry into cells. In this context an analysis via the integrated gradients method shows that only parts of the sigma 2 receptor and the SARS-CoV-2 spike protein contribute to our model’s prediction of the protein-protein interaction (Figure S6). More specifically, the non-cytoplasmic domain and EXPERA domains demonstrate positive integrated gradient scores, i.e. the existence of these domains influence our model to make the according prediction.

## Conclusion

Huge advancements have been made recently by applying deep learning algorithms from the natural language processing field to protein bioinformatics. Protein language models such as ProtTrans and ProtBERT (Elnaggar et al. 2021) trained on billions of protein sequences learn informative features through the transformation of sequences to vector representations. These models previously showed their predictive power in various tasks such as prediction of secondary structure or classification of membrane proteins (Elnaggar et al. 2021).

In our work, we used ProtBERT within a specifically designed Siamese neural network architecture to predict PPIs by only utilizing the primary sequences of protein pairs. We trained our models following a PU learning scheme and performed an extensive evaluation and hyperparameter optimization of our models, demonstrating high prediction performances for virus protein to human receptor interactions of JCV and SARS-CoV-2. An additional head to head comparison to the recently-published method by Tsukiyama et al. (2021) using a more traditional word2vec sequence embedding combined with an LSTM unit revealed state-of-the art prediction performance of our STEP approach.

Interactions predicted by our proposed model between JCV major capsid protein VP1 and receptors in brain cells showed a strong enrichment of different neurotransmitters, including serotonin receptors, which is in line with the current literature. For the SARS-Cov-2 spike protein, our model interestingly predicted for all virus variants an interaction with the sigma intracellular receptor 2, which might explain the cytopathic effects of sigma receptor binding ligands reported in the literature (Abate et al. 2020, Gordon et al. 2020, Ostrov et al. 2021). In both cases recent techniques coming from the field of Explainable AI (XAI) allowed us to interpret model predictions and identify those parts of protein sequences, which according to our model mostly influence the prediction of respective protein-protein interactions. Of course, a validation of these predictions would require experimental procedures, which are beyond the scope of this paper.

Altogether, our work demonstrates the potential of modern deep learning-based biological sequence embeddings and modern XAI techniques for bioinformatics. While in this paper we focused on John Cunningham polyomavirus and SARS-CoV-2, our proposed model could in future work be easily trained to predict interactions of other viruses as well and thus contribute to the emerging set of computational methods that might help to respond to future epidemic and pandemic situations more effectively. In addition, there is the potential to use our method in the context of modern drug development approaches, which employ virus-like particles to deliver compounds to specific tissues and receptors. We have made our method publically available via github.

## Acknowledgements

We thank André Gemünd for his support regarding the computational infrastructure of SCAI.

## Author Contributions

Conceptualization, O.E. and H.F.; Methodology, H.F. and S.M.; Data Curation, Formal Analysis, Visualization, Investigation, Validation, S.M.; Supervision, H.F.; Project Administration, V.D., M.S., O.E., and H.F.; Writing -Original Draft, S.M. and H.F.; Writing - Review & Editing, S.M., V.D., M.S., O.E., and H.F.

