## Supplementary Material for "Accurate Prediction of Virus-Host Protein-Protein Interactions via a Siamese Neural Network Using Deep Protein Sequence Embeddings"

#### Additional Files

File S1. Includes brain-specific tissue types (is referenced in Section "Construction of Brain-specific PPI Dataset")
File S2. Includes all identified human receptors
File S3. Includes prediction results of John Cunningham polyomavirus major capsid protein VP1 (https://www.uniprot.org/uniprot/P03089, accessed on 18 November 2021)
File S4. Includes prediction results of omicron variant of SARS-CoV-2 spike protein (https://www.uniprot.org/uniprot/P0DTC2, accessed on 18 November 2021)

#### Comparative Evaluation of STEP with state-of-the-art Work

We performed a head-to-head comparison of our STEP architecture on three different PPI detection datasets published by Tsukiyama et al. (2021), Guo et al. (2008), and Sun et al. (2017). Tables S1, S2, and S3 show the detailed results of these experiments.

Table S1: Overview of the results of comparative evaluation of STEP with LSTM-PHV (Tsukiyama et al. 2021) work. Similarly, as the authors of LSTM-PHV, we applied a 5-fold cross validation using the published dataset for training the STEP models. The highest values are highlighted in bold.

| | LSTM-PHV (Tsukiyama et al. 2021) | | STEP (ours) | |
|---|---|---|---|---|
| | **AUC** | **AUPR** | **AUC** | **AUPR** |
| Fold 1 | 97.60% | 94.10% | **98.59%** | **95.01%** |
| Fold 2 | 97.80% | 94.40% | **98.97%** | **96.22%** |
| Fold 3 | 97.60% | 93.40% | **98.51%** | **95.32%** |
| Fold 4 | 97.50% | 93.70% | **98.70%** | **95.63%** |
| Fold 5 | 97.40% | 93.70% | **98.82%** | **96.35%** |
| Average over 5 folds | 97.60% | 93.90% | **98.72% (+/-0.16%)** | **95.71% (+/-0.51%)** |

Table S2: Overview of the results of comparative evaluation of STEP on Yeast dataset (Guo et al. 2008) work. As Chen et al. (2019), we also applied a 5-fold cross validation using the dataset for training the STEP models.

| | **AUC** | **AUPR** | **F1** | **MCC** |
|---|---|---|---|---|
| Fold 1 | 99.74% | 99.75% | 97.47% | 95.02% |
| Fold 2 | 99.47% | 99.30% | 97.36% | 94.73% |
| Fold 3 | 99.69% | 99.72% | 97.62% | 95.27% |
| Fold 4 | 99.63% | 99.65% | 97.53% | 95.09% |
| Fold 5 | 99.51% | 99.49% | 96.85% | 93.75% |
| Average over 5 folds | 99.61% (+/-0.10%) | 99.58% (+/-0.17%) | 97.37% (+/-0.27%) | 94.77% (+/-0.54%) |

Table S3: Overview of the results of comparative evaluation of STEP on Human dataset (Sun et al. 2017) work. As Sun et al. 2017, we also applied a 10-fold cross validation using the dataset for training the STEP models.

| | **AUC** | **AUPR** | **F1** | **MCC** |
|---|---|---|---|---|
| Fold 1 | 99.76% | 99.65% | 98.89% | 97.78% |
| Fold 2 | 99.74% | 99.68% | 98.85% | 97.70% |
| Fold 3 | 99.74% | 99.70% | 98.79% | 97.60% |
| Fold 4 | 99.78% | 99.73% | 98.84% | 97.68% |
| Fold 5 | 99.72% | 99.61% | 98.71% | 97.43% |
| Fold 6 | 99.74% | 99.66% | 98.70% | 97.40% |
| Fold 7 | 99.71% | 99.68% | 98.76% | 97.51% |

| | | | | |
|---|---|---|---|---|
| Fold 8 | 99.67% | 99.59% | 98.95% | 97.89% |
| Fold 9 | 99.72% | 99.69% | 98.97% | 97.95% |
| Fold 10 | 99.77% | 99.63% | 98.89% | 97.78% |
| Average over 10 folds | 99.74% (+/-0.03%) | 99.66% (+/-0.04%) | 98.84% (+/-0.09%) | 97.67% (+/-0.18%) |

#### Interaction Type Prediction and Binding Affinity Estimation

We also evaluated our STEP architecture on two different tasks, namely PPI type prediction and a PPI binding affinity estimation. For PPI type prediction, we used the SHS27k dataset that was extracted by Chen et al. (2019) from the STRING database. It contains 26,944 instances that are distributed among the seven interaction types: activation (16.70%), binding (16.70%), catalysis (16.70%), expression (5.84%), inhibition (16.70%), post-translational modification (ptmod; 10.66%), and reaction (16.70%). The dataset for PPI binding affinity estimation was collected from SKEMPI (the structural database of kinetics and energetics of mutant protein interactions) database. The final dataset contains the binding affinity of 2,792 mutant protein complexes. We refer to Chen et al. (2019) for the details on the construction of this dataset.

The STEP architecture was slightly modified for the two tasks. The PPI type prediction was considered as a multi-class prediction problem. We replaced our bottleneck classification head (see Figure 1) with three identical linear layers (in addition to dropout and ReLU activation layers). The loss function was replaced with cross-entropy loss and the sigmoid activation function was replaced with Softmax to cope with the given seven interaction type classes. For the PPI binding affinity estimation task, which is considered a regression problem, we just replaced the loss function with mean squared error loss as Chen et al. (2019). Chen et al. (2019) also applied a min-max scaling with range between 0 and 1 on the binding affinity value in the training set; we used the same transformation. The experiments for both tasks were performed in a 10-fold CV setting using the same data splits as Chen et al. (2019). Table S4 contains the results of both experiments, which shows that STEP with the usage of transfer learning reaches state-of-the-art results on both datasets.

Table S4. Overview of the results of comparative evaluation of STEP on PPI interaction type prediction and binding affinity estimation tasks (Chen et al. 2019). For the PPI interaction type prediction, we report the zero rule baseline and accuracy the fold changes over zero rule. For the STEP model, we also report macro $F_1$ and MCC scores. For the PPI binding affinity task, we report mean squared error, mean absolute error, and the Pearson correlation coefficient. As Chen et al. (2019), we also applied a 10-fold cross validation using the dataset for training the STEP models. The highest values are highlighted in bold. MCC = Matthews correlation coefficient MSE = mean squared error; MAE = mean absolute error.

**Comparative analysis on PPI type prediction dataset (SHS27k) from Chen et al. (2019) via 10-fold cross-validation**

| | **Accuracy (foldx)** | **$F_1$ (macro)** | **MCC** |
|---|---|---|---|
| **Zero rule (baseline)** | 16.70% (1.00x) | - | - |
| **Chen et al. (2019)** | 59.56% (3.57x) | NA | NA |
| **STEP (ours)** | **59.77% (3.58x)** | 59.85% (+/- 1.10%) | 54.87% (+/-1.13%) |

**Comparative analysis on PPI binding affinity estimation dataset from Chen et al. (2019) via 10-fold cross-valiation**

| | **MSE (x $10^{-2}$)** | **MAE (x $10^{-2}$)** | **Pearson Corr. Coef.** |
|---|---|---|---|
| **Chen et al. (2019)** | 0.63 | 5.48 | 0.873 |
| **STEP (ours)** | **0.2685** | **3.5214** | **0.9413** |

### Prediction of JCV Major Capsid Protein VP1 Interactions

#### Overview on the frequency of interactions for brain-specific tissue types

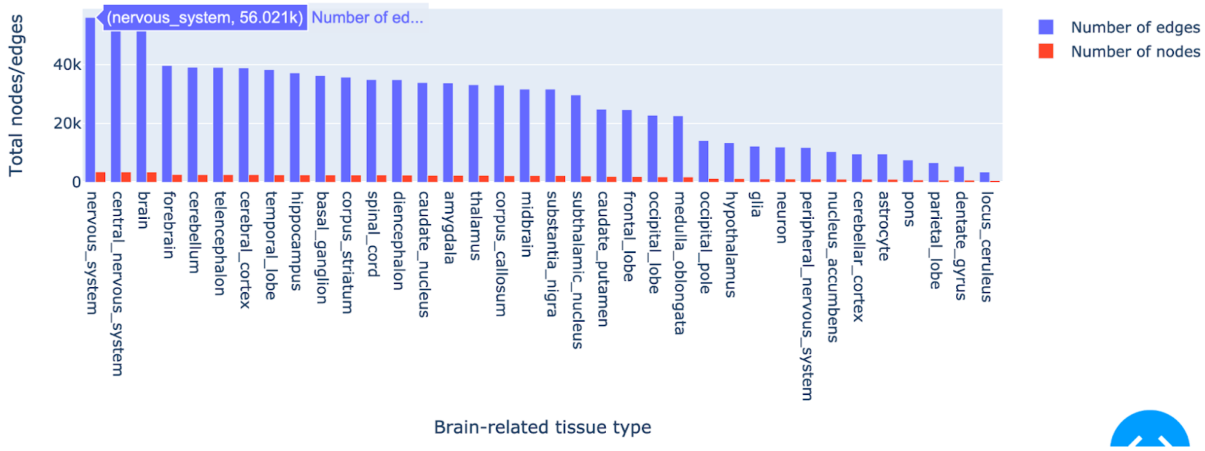

Figure S1. Overview on the frequency of interactions for brain-specific tissue types

#### Hyperparameters of STEP-Brain Model

Table S5: Overview of STEP-Brain model hyperparameters tuned with Bayesian hyperparameter optimization. Hyperparameters were sampled from continuous distributions (such as Adam epsilon, weight decay) and categorical (such as accumulate grad batches) spaces.

| **Hyperparameter** | **Search range** | **Final model values** |
|---|---|---|
| Dense layer 1 | fix | 4096 (in), 256 (out) |
| Dense layer 2 | fix | 256 (in), 16 (out) |
| Dense layer 3 | fix | 16 (in), 1 (out) |
| Adam epsilon | [1e-10, 1e-6] | 2.6315108252722756e-10 |
| Weight decay | [1e-10, 1e-1] | 1.4955169166544313e-06 |
| Accumulate grad batches | {16, 32, 64, 128} | 32 |
| Dropout layer 1 | [0.2, 0.5] | 0.3326435551602387 (prob), 4096 (in), 4096 (out) |
| Dropout layer 2 | fix | 0.2 (prob), 256 (in), 256 (out) |
| Dropout layer 3 | fix | 0.2 (prob), 16 (in), 16 (out) |
| Number of frozen epochs | [1, 5] | 3 |
| learning rate | [1e-05, 0.005] | 7.091867154042537e-05 |
| optimizer | fix | AdamW |

#### Results on Extended Test Dataset

Our test set consists of an equal number of positive and pseudo-negative samples. To investigate the effect of an increased number of pseudo-negative instances in the test set, we extended it with random samples so that the ratio of positive to pseudo-negative samples is 1:10. In Table S6 we report the results for the both test sets. The results show that the model performance stays stable for both ratios.

Table S6: Overview of performance of the STEP-Brain model on the test set with equal number of pseudo-negatives and true positive PPIs (1:1) and 10 times more pseudo-negatives than true positive PPIs (1:10).

| **Positive to pseudo-negative ratio in test set** | **AUC** | **AUPR** |
|---|---|---|
| 1:1 | 88.78% | 88.32% |

| | | |
|---|---|---|
| 1:10 | 89.00% | 98.59% |

#### Significant GO Molecular Functions and Enrichment Map of the Capsid Protein VP1 Interactions

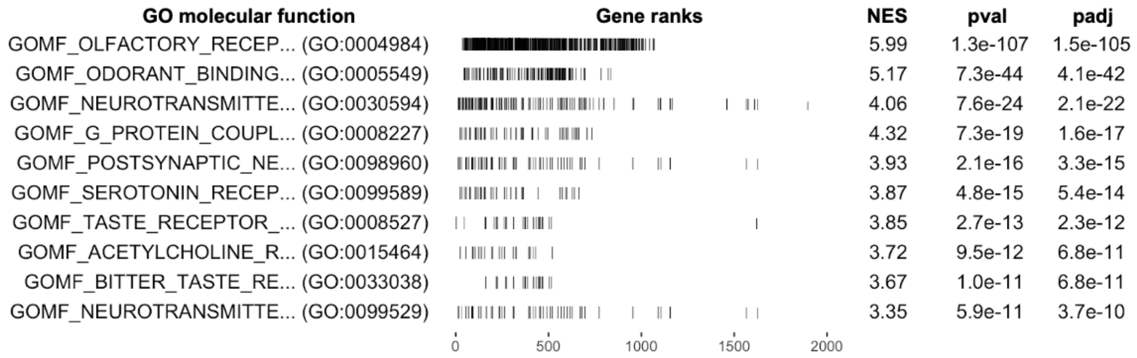


Figure S2. Top 10 significant GO molecular functions retrieved by GSEA for the predicted capsid protein VP1 and human receptor interactions. Column GO molecular function contains the name and the ID of the GO molecular function, Gene ranks shows the distribution of significant genes. NES stands for normalized enrichment score, pval is the original p-value, and padj is the false discovery rate adjusted p-value.

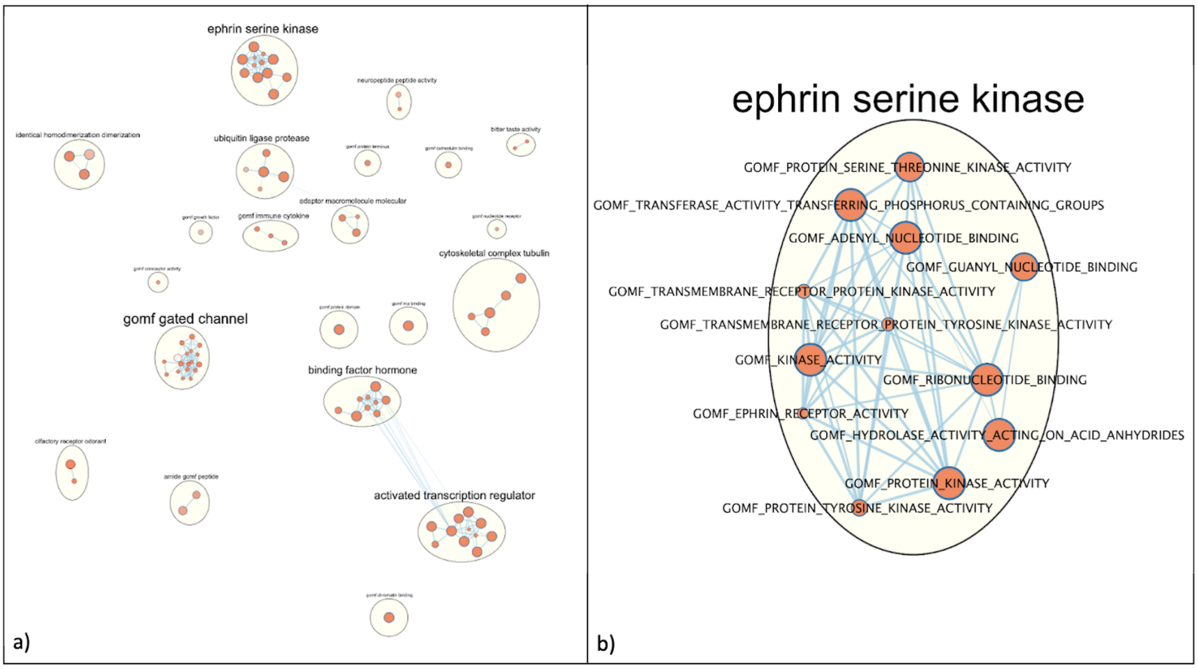


Figure S3. a) Enrichment map of significant GO molecular functions retrieved by GSEA for the predicted capsid protein VP1 and human receptor interactions. b) a detailed version of “ephrin serine kinase” cluster with the included GO molecular functions.

#### Integrated Gradients Analysis for JCV Capsid VP1 Protein

In this analysis, we use the integrated gradients (IG) method (Sundararajan et al. 2017) to interpret the predictions made by our STEP-Brain model and visualize attributions of each of the amino acids. Also, we annotate the sequence regions with information (such as domains and structural features) from the InterPro database. Figure S4 shows the IG scores of each amino acid for the interaction between UPF0606 protein KIAA1549 (top) and JCV major capsid VP1 protein (bottom). The visualization includes the IG (attribution) score and InterPro annotations of both sequences. The overall attribution score defines the sum of all amino acid attribution scores based on the contribution of each amino acid to the model's output. The amino acid importance highlights each amino acid of the given sequence and shows its positive, negative, or neutral contribution to the predicted class. Here, both sequences have multiple regions that have high positive attribution scores. For the UPF0606 protein KIAA1549 (top) sequence, various structural features could be obtained through the InterPro database. Signal peptide N-regions negatively contribute to the predicted class, whereas the beginning of the non-cytoplasmic domain region is contributing positively. Further regions in the samedomain contribute negatively. Unfortunately, no structural and domain information is available for the JCV major capsid VP1 protein (bottom).

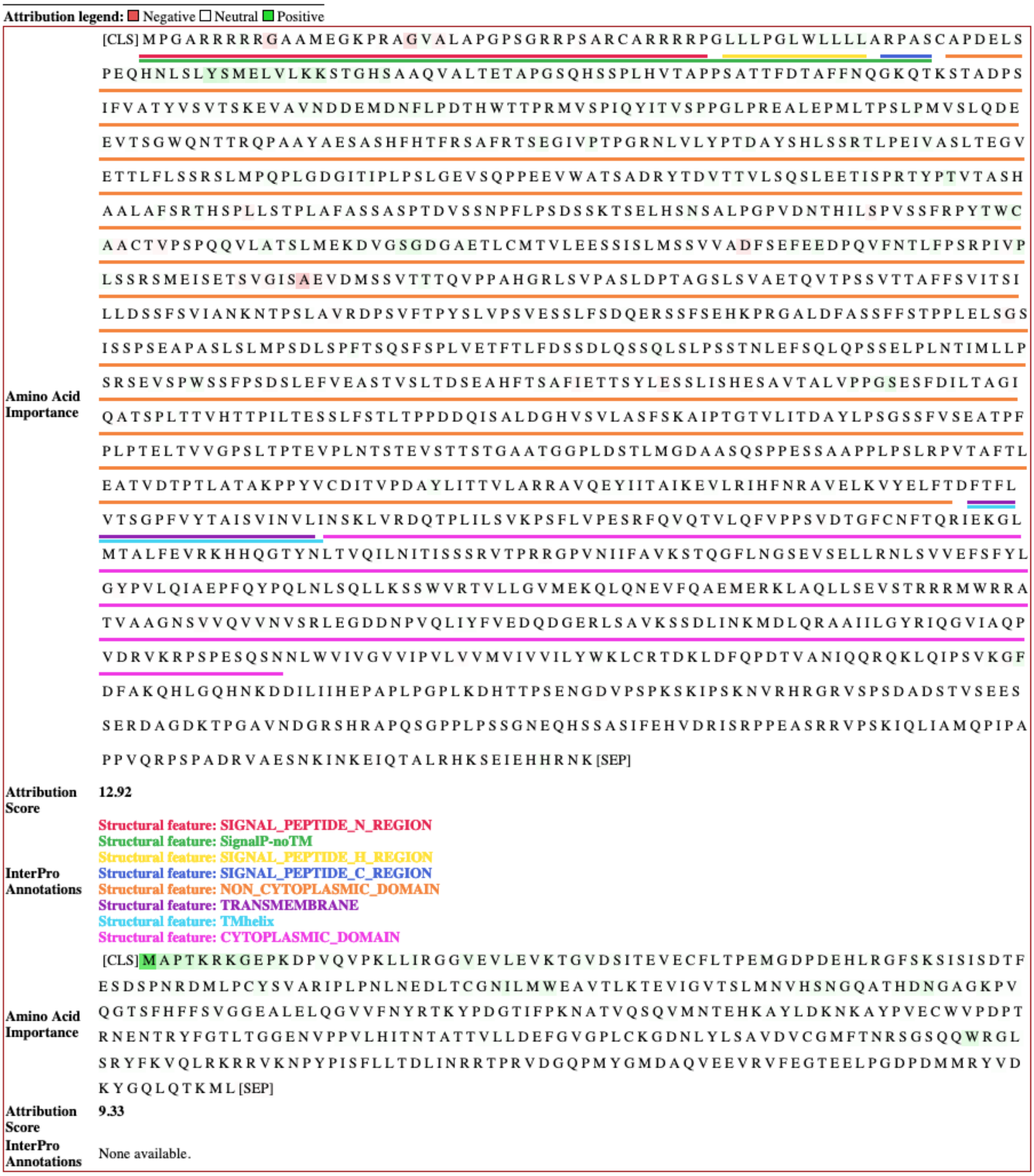


Figure S4. Visualization of the attributions of each amino acid for the interaction between UPF0606 protein KIAA1549 (top) and JCV major capsid VP1 protein (bottom).

#### InterPro Domain Enrichment Analysis

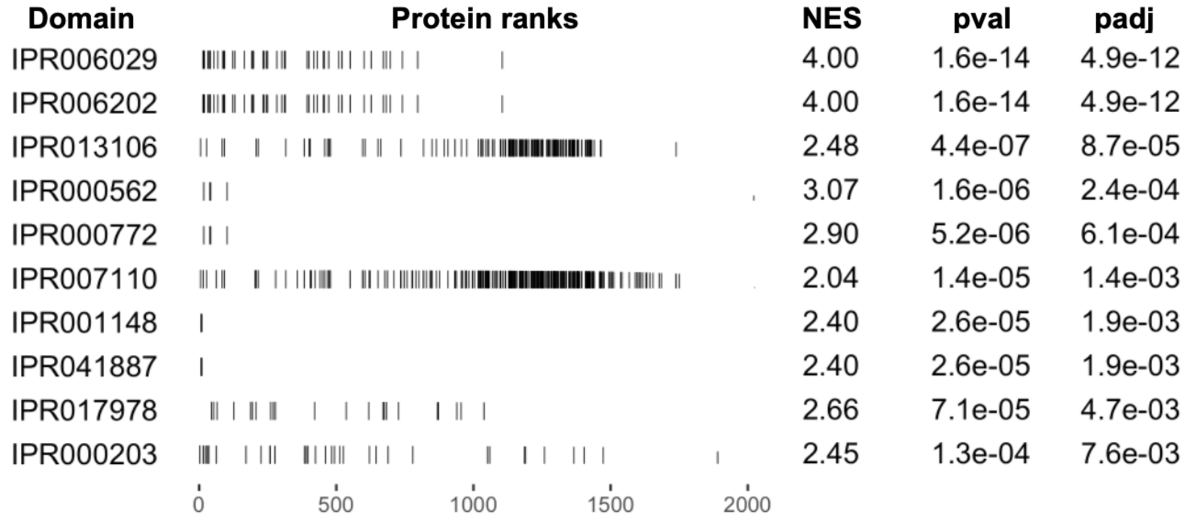


Figure S5. Top 10 significant InterPro domains retrieved by enrichment analysis for the predicted capsid protein VP1 and human receptor interactions. Column Domain contains the ID of the InterPro domain, Protein ranks shows the distribution of significant proteins. NES stands for normalized enrichment score, pval is the original p-value, and padj is the false discovery rate adjusted p-value.

Table S7. ID, name, and weblink of top 10 significant InterPro domains retrieved by the enrichment analysis (see Figure S5).

| InterPro ID | Name | Weblink |
|---|---|---|
| IPR006029 | Neurotransmitter-gated ion-channel transmembrane domain | https://www.ebi.ac.uk/interpro/entry/InterPro/IPR006029/ |
| IPR006202 | Neurotransmitter-gated ion-channel ligand-binding domain | https://www.ebi.ac.uk/interpro/entry/InterPro/IPR006202/ |
| IPR013106 | Immunoglobulin V-set domain | https://www.ebi.ac.uk/interpro/entry/InterPro/IPR013106/ |
| IPR00562 | Fibronectin type II domain | https://www.ebi.ac.uk/interpro/entry/InterPro/IPR000562/ |
| IPR00772 | Ricin B, lectin domain | https://www.ebi.ac.uk/interpro/entry/InterPro/IPR000772/ |
| IPR007110 | Immunoglobulin-like domain | https://www.ebi.ac.uk/interpro/entry/InterPro/IPR007110/ |
| IPR001148 | Alpha carbonic anhydrase domain | https://www.ebi.ac.uk/interpro/entry/InterPro/IPR001148/ |
| IPR041887 | Receptor-type tyrosine-protein phosphatase, carbonic anhydrase domain | https://www.ebi.ac.uk/interpro/entry/InterPro/IPR041887/ |

| | | |
|---|---|---|
| IPR017978 | GPCR family 3, C-termina | https://www.ebi.ac.uk/interpro/entry/InterPro/IPR017978/ |
| IPR000203 | GPS motif | https://www.ebi.ac.uk/interpro/entry/InterPro/IPR000203/ |

### Prediction of SARS-CoV-2 Spike Glycoprotein Interactions

#### Hyperparameters of STEP-Virus-Host Model

Table S8: Overview of STEP-Virus-Host model hyperparameters tuned with Bayesian hyperparameter optimization. Hyperparameters were sampled from continuous distributions (such as Adam epsilon, weight decay) and categorical (such as accumulate grad batches) spaces.

| **Hyperparameter** | **Search range** | **Final model values** |
|---|---|---|
| Dense layer 1 | fix | 4096 (in), 256 (out) |
| Dense layer 2 | fix | 256 (in), 16 (out) |
| Dense layer 3 | fix | 16 (in), 1 (out) |
| Adam epsilon | [1e-10, 1e-6] | 5.90539e-08 |
| Weight decay | [1e-10, 1e-1] | 6.34672e-07 |
| Accumulate grad batches | {16, 32, 64, 128} | 32 |
| Dropout layer 1 | [0.2, 0.5] | 0.4 (prob), 4096 (in), 4096 (out) |
| Dropout layer 2 | fix | 0.2 (prob), 256 (in), 256 (out) |
| Dropout layer 3 | fix | 0.2 (prob), 16 (in), 16 (out) |
| Number of frozen epochs | [1, 5] | 4 |
| learning rate | [1e-05, 0.005] | 0.000429331 |
| optimizer | fix | AdamW |

#### Results on Extended Test Dataset

Similar to the first use case, we also investigated the effect of an increased number of pseudo-negative instances in the test set, by extending it with random samples so that the ratio of positive to pseudo-negative samples is 1:10. In Table S9 we report the results for the both test sets. Detailed results are reported in Table S10. The results show that AUC is stable for both test sets but AUPR falls significantly.

Table S9: Overview of performance of the STEP-Virus-Host model on the test set with equal number of pseudo-negatives and true positive PPIs (1:1) and 10 times more pseudo-negatives than true positive PPIs (1:10).

| Positive to pseudo-negative ratio in test set | AUC | AUPR |
|---|---|---|
| 1:1 | 83.42% (+/- 3.91%) | 84.02% (+/- 4.58%) |
| 1:10 | 84.65% (+/- 0.90%) | 41.90% (+/-5.07%) |

#### Additional Nested Cross-Validation Results

Table S10: Results of the outer loop folds retrieved during the nested cross validation of STEP-Virus-Host model by using the test set with an ratio of 1:10 positive to pseudo-negative instances.

| Outer Fold | AUC | AUPR |
|---|---|---|
| 1 | 85.14% | 43.99% |
| 2 | 85.49% | 50.12% |
| 3 | 83.01% | 36.17% |
| 4 | 85.22% | 37.01% |
| 5 | 84.38% | 42.21% |
| **Mean** | 84.65% (+/- 0.90%) | 41.90% (+/- 5.07%) |

#### Predictions of Spike Glycoprotein of SARS-CoV-2 Omicron, Alpha, and Delta Variants

Table S11: Top 10 predicted interactions of the spike protein of omicron variant of SARS-CoV-2 and human receptors ranked by the probability score obtained by our model.

| Rank | Receptor protein ID | Receptor protein name | Score (in %) |
|---|---|---|---|
| 1 | Q5BJF2 | Sigma intracellular receptor 2 | 73.14% |
| 2 | Q8WWA0 | Intelectin-1 | 17.02% |
| 3 | P55085 | Proteinase-activated receptor 2 | 16.16% |
| 4 | Q15546 | Monocyte to macrophage differentiation factor | 13.94% |
| 5 | O15243 | Leptin receptor gene-related protein | 13.64% |
| 6 | O95214 | Leptin receptor overlapping transcript-like 1 | 13.47% |
| 7 | Q9Y2W1 | Thyroid hormone receptor-associated protein 3 | 13.38% |

| | | | |
|---|---|---|---|
| 8 | Q96Q45 | Transmembrane protein 237 | 13.37% |
| 9 | P0DPR3 | T cell receptor delta diversity 1 | 12.71% |
| 10 | Q9UJM3 | ERBB receptor feedback inhibitor 1 | 12.44% |

Table S12: Top 10 predicted interactions of the spike protein of alpha variant of SARS-CoV-2 and human receptors ranked by the probability score obtained by our model.

| Rank | Receptor protein ID | Receptor protein name | Score (in %) |
|---|---|---|---|
| 1 | Q5BJF2 | Sigma intracellular receptor 2 | 71.82% |
| 2 | Q8WWA0 | Intelectin-1 | 18.96% |
| 3 | P55085 | Proteinase-activated receptor 2 | 17.03% |
| 4 | O15243 | Leptin receptor gene-related protein | 14.03% |
| 5 | O95214 | Leptin receptor overlapping transcript-like 1 | 13.77% |
| 6 | Q15546 | Monocyte to macrophage differentiation factor | 13.52% |
| 7 | Q96Q45 | Transmembrane protein 237 | 13.04% |
| 8 | P0DPR3 | T cell receptor delta diversity 1 | 12.71% |
| 9 | P60201 | Myelin proteolipid protein | 12.30% |
| 10 | Q9Y6Y9 | Lymphocyte antigen 96 | 12.17% |

Table S13: Top 10 predicted interactions of the spike protein of delta variant of SARS-CoV-2 and human receptors ranked by the probability score obtained by our model.

| Rank | Receptor protein ID | Receptor protein name | Score (in %) |
|---|---|---|---|
| 1 | Q5BJF2 | Sigma intracellular receptor 2 | 73.87% |
| 2 | Q8WWA0 | Intelectin-1 | 18.47% |
| 3 | Q15546 | Monocyte to macrophage differentiation factor | 15.10% |
| 4 | P55085 | Proteinase-activated receptor 2 | 14.93% |
| 5 | O15243 | Leptin receptor gene-related protein | 14.43% |
| 6 | O95214 | Leptin receptor overlapping transcript-like 1 | 14.31% |

| | | | |
|---|---|---|---|
| 7 | Q96Q45 | Transmembrane protein 237 | 14.02% |
| 8 | P0DPR3 | T cell receptor delta diversity 1 | 13.38% |
| 9 | Q9UJM3 | ERBB receptor feedback inhibitor 1 | 12.93% |
| 10 | P60201 | Myelin proteolipid protein | 12.60% |

#### Integrated Gradients Analysis for SARS-CoV-2 Spike Glycoprotein

Here, we use the integrated gradients method (Sundararajan et al. 2017) to interpret the predictions made by our STEP-Virus-Host model and visualize IG scores (attributions) of each of the amino acids. Also, we annotate the sequence regions with information (such as domains and structural features) from the InterPro database (Blum et al. 2021). Figure S6 shows the attributions of each amino acid for the interaction between Sigma intracellular receptor 2 (top) and omicron variant of SARS-CoV-2 Spike Glycoprotein (bottom). The visualization includes the amino acid importance, the attribution score, and InterPro annotations of both sequences. The overall attribution score defines the sum of all amino acid attribution scores based on the contribution of each amino acid to the model's output. The amino acid importance highlights each amino acid of the given sequence and shows its positive, negative, or neutral contribution to the predicted class. In this case the attribution score of the Sigma intracellular receptor 2 (top) sequence is positive, showing that the majority of the amino acids have positively contributed to the predicted class. The cytoplasmic domain negatively contributes to the interaction, whereas the contribution of the non-cytoplasmic domain and EXPERA domain is positive. The attribution score of SARS-CoV-2 Spike Glycoprotein (bottom) is negative. It is worthy to note that the attributing regions are located mostly in the "S1 subunit, receptor-binding domain" (InterPro:IPR018548), which is also known for binding to angiotensin-converting enzyme 2 (ACE2) in respiratory syndrome coronavirus (SARS-CoV) (Prabakaran et al. 2006, Journal of Biological Chemistry).

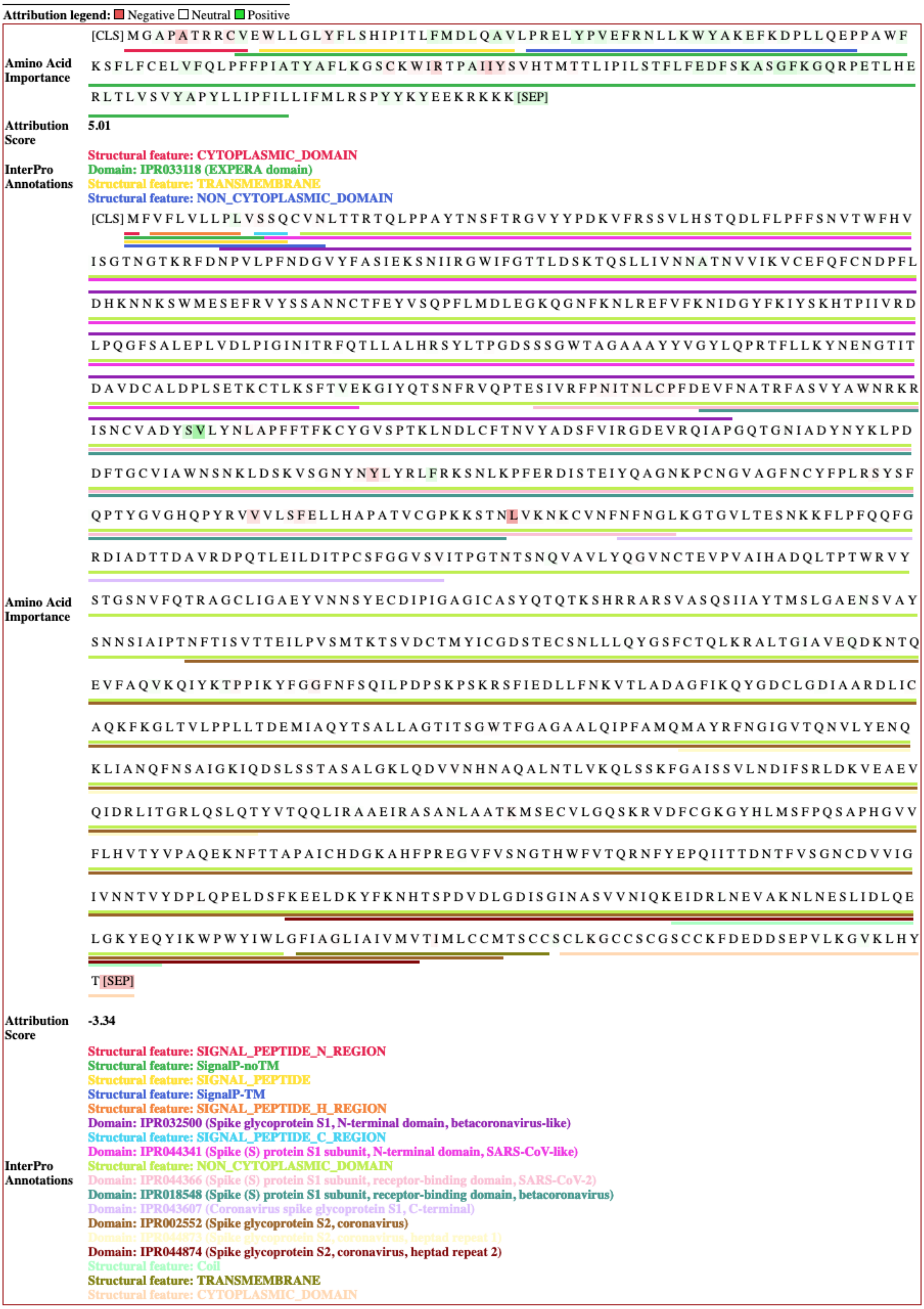


Figure S6. Visualization of the attributions of each amino acid for the interaction between Sigma intracellular receptor 2 (top) and omicron variant of SARS-CoV-2 Spike Glycoprotein (bottom).

#### Implementation Details

The implementation of the introduced work is performed in Python (Van Rossum et al. 2009). For the exploratory data analysis and data processing we used Pandas (The pandas development team 2020, Wes McKinney 2010), Sqlite3 (https://www.sqlite.org/, accessed on 18 November 2021), NetworkX (https://networkx.org/, accessed on 18 November 2021) and Plotly (https://plotly.com/, accessed on 18 November 2021). We used the frameworks PyTorch (https://pytorch.org/, accessed on 18 November 2021), PyTorch Lightning (https://www.pytorchlightning.ai/, accessed on 18 November 2021) and Transformers (https://huggingface.co/transformers/, accessed on 18 November 2021) for the implementation of the models. The metrics were calculated using the TorchMetrics (https://torchmetrics.readthedocs.io/en/latest/, accessed on 18 November 2021) and scikit-learn (https://scikit-learn.org/stable/, accessed on 18 November 2021) libraries. All experiments were tracked via MLflow (https://www.mlflow.org/, accessed on 18 November 2021). The Bayesian hyperparameter optimization was performed with Optuna (Akiba et al. 2019) (https://optuna.readthedocs.io/en/stable/, accessed on 18 November 2021). MariaDB (https://mariadb.org/, accessed on 18 November 2021) was used by Optuna and MLFlow to store intermediate and final results of experiments. GSEA was performed using the R library fgsea (https://bioconductor.org/packages/release/bioc/html/fgsea.html, accessed on 18 November 2021). Furthermore, the library Captum (https://captum.ai/, accessed on 24 February 2022) was used to calculate integrated gradients. The HPC Infrastructure (https://www.scai.fraunhofer.de/en/about-us/hpc-center.html, accessed on 18 November 2021) of Fraunhofer Institute SCAI that includes the GPU compute nodes was accessed via Slurm (https://slurm.schedmd.com/, accessed on 18 November 2021).
